# Genetic control of a sex-specific piRNA program

**DOI:** 10.1101/2022.10.25.513766

**Authors:** Peiwei Chen, Alexei A. Aravin

**Author notes:** To whom correspondence should be addressed: Alexei A. Aravin, Peiwei Chen.

## Abstract

Sexually dimorphic traits in morphologies are widely studied, but those in essential molecular pathways remain largely unexplored. Previous work showed substantial sex differences in *Drosophila* gonadal piRNAs, which guide PIWI proteins to silence selfish genetic elements thereby safeguarding fertility. However, the genetic control mechanisms of piRNA sexual dimorphism remain unknown. Here, we showed that most sex differences in the piRNA program originate from the germline rather than gonadal somatic cells. Building on this, we dissected the contribution of sex chromosome and cellular sexual identity towards the sex-specific germline piRNA program. We found that the presence of the Y chromosome is sufficient to recapitulate some aspects of the male piRNA program in a female cellular environment. Meanwhile, sexual identity controls the sexually divergent piRNA production from X-linked and autosomal loci, revealing a crucial input from sex determination into piRNA biogenesis. Sexual identity regulates piRNA biogenesis through Sxl and this effect is mediated in part through chromatin proteins Phf7 and Kipferl. Together, our work delineated the genetic control of a sex-specific piRNA program, where sex chromosome and sexual identity collectively sculpt an essential molecular trait.

## Main

Sexual dimorphism, where a trait is modified by the biological sex to manifest in distinct ways between males and females, is pervasive in nature. While sexually dimorphic traits in morphologies have been widely studied^1–4^, those in essential molecular pathways remain largely unexplored. In *Drosophila melanogaster* gonads, the piRNA program executes a critical function by guiding the PIWI-clade Argonaute proteins to silence selfish genetic elements such as transposons^5–7^, thereby safeguarding fertility. To pass on the transgenerational memory of proper piRNA targets, mothers deposit piRNAs to the embryo, instructing the zygotic genome to mount a homologous piRNA program in the next generation that reflects the maternal response to genomic parasites^8–11^. However, males implement a piRNA program distinct from their female siblings^12^, the underlying mechanism of which is elusive. We previously found evidence for both differential transcription of piRNA loci in the nucleus^13^ and differential processing of piRNA precursor transcripts in the cytoplasm^12^ between the two sexes, but the upstream control of these sexually dimorphic molecular events is unknown. In this work, we sought to decipher the genetic control of piRNA sexual dimorphism, in order to gain insights into the mechanisms by which sexual dimorphism in essential molecular traits is sculpted.

## Results

Prior work compared the male and female piRNA profiles from two different *D. melanogaster* lab strains^12^, where distinct genetic backgrounds confounded the characterization of piRNA sexual dimorphism. In addition, the sex of *D. melanogaster* is determined independently of the presence of the Y chromosome (both XY and XO flies are phenotypic males, while XX and XXY flies are phenotypic females)^14–16^, so the morphology-based identification of males and females does not directly translate to an interpretation of Y chromosome status. Given that several piRNA-producing loci reside on the Y^12^, the inability to infer Y chromosome content from the phenotypic sex complicates the characterization of piRNA sexual dimorphism. To circumvent these issues, we introduced a Y chromosome marked by *y*^+^ and *w*^+^ genes (hereafter *y^+^w^+^*Y) into an inbred *yw* stock and backcrossed it to *yw* for multiple consecutive generations (see methods). This line allowed us to unequivocally identify XY males (red-eyed flies with black body color and male genitalia) and XX females (white-eyed flies with yellow body color and female genitalia) (Fig. 1a), from which we profiled the gonadal piRNAs in each of the two sexes. Analysis of the piRNA libraries showed substantial intersexual differences in the abundance of piRNAs targeting different transposon families (Fig. 1b) and expression levels of individual major piRNA loci in the genome (Fig. 1c), largely in agreement with our previous study^12^. Having excluded the possible confounding effects of genetic backgrounds and Y chromosome status, we confirmed that the piRNA program in *D. melanogaster* gonads is sexually dimorphic.

**Fig. 1.**
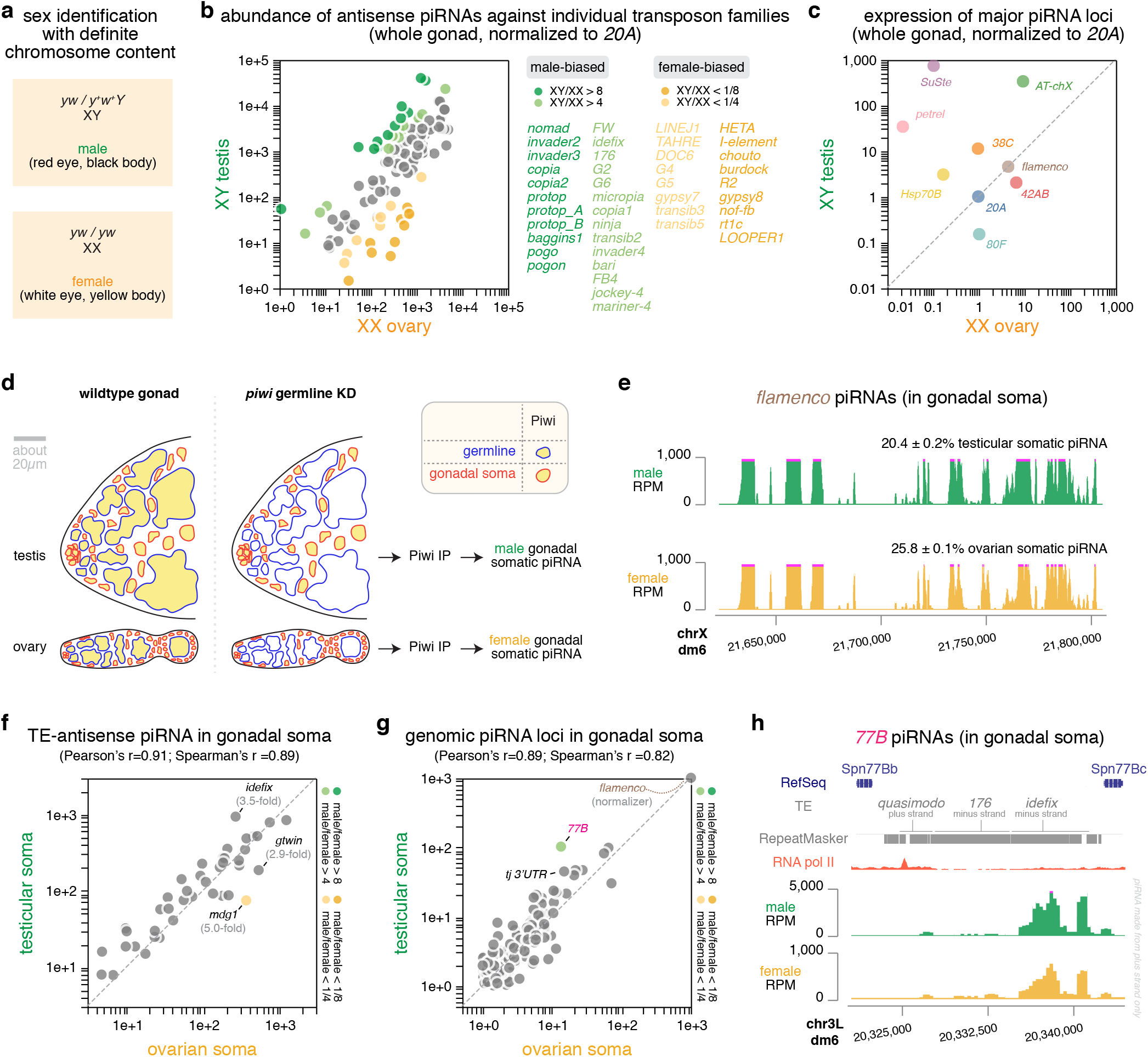
Germline is the major cell type origin of piRNA sexual dimorphism in *D. mel* gonads. **a**, Genotype and phenotype of males and females that can be identified with definite chromosome content, employing an X chromosome lacking *y* and *w* genes as well as a Y chromosome that carries the wildtype *y*^+^ and *w*^+^ genes. **b**, Comparison of the abundance of piRNAs targeting individual transposon families in XY testis versus XX ovary, normalized to the expression of *20A* piRNAs. Sex-biased transposon-targeting piRNAs are color coded and listed on the right. **c**, Comparison of the expression of major piRNA loci in the genome^12^ in XY testis versus XX ovary, normalized to the expression of *20A* piRNAs. Each locus is marked by a different color. **d**, Illustration of the experimental strategy to isolate somatic piRNAs in the gonad. Left: cartoon showing the cell type composition of testis and ovary, with germline having a blue outline and gonadal somatic cells having a red outline. Both germline and gonadal somatic cells express Piwi, which is marked by yellow. Right: cartoon showing Piwi expression in testis and ovary upon efficient, germline-specific knock-down of *piwi* that completely depletes Piwi in the germline, leaving the somatic cells as the only source of Piwi in the gonad. Gonadal somatic piRNAs are isolated by immunoprecipitating Piwi from these gonads that lose germline Piwi. **e**, UCSC genome browser view of the *flamenco* piRNA locus showing *flamenco* piRNAs take up similar fractions of gonadal somatic piRNAs in testis and ovary with similar coverage profiles. **f**, Comparison of the abundance of transposon-antisense piRNAs in testicular and ovarian soma, normalized to the expression of *flamenco* piRNAs. Sex-biased piRNAs are color coded in the same way as in **b** and the correlation coefficients are reported. **g**, Comparison of the expression of different piRNA loci in testicular and ovarian soma, normalized to the expression of *flamenco* piRNAs. Sex-biased piRNAs are color coded in the same way as in **b** and the correlation coefficients are reported. **h**, UCSC genome browser view of the *77B* piRNA locus, showing its flanking protein-coding genes, its transposon contents and piRNA coverage profiles in two sexes. Note the difference in y-axis scales that reflects a higher relative activity of *77B* in the testicular soma than the female counterpart. A putative promoter marked by an RNA pol II peak likely drives the expression of piRNAs from the plus strand that are antisense to two transposons*, 176* and *idefix*.

### Germline is the major cell type origin of piRNA sexual dimorphism

In *D. melanogaster* gonads, the piRNA program operates in both the germline and gonadal somatic cells, but piRNA biogenesis and targets differ between the two cell types^9,17^. Thus, the male-female differences seen in gonad-wide piRNA quantification could reflect sexual dimorphism in either germline or gonadal somatic cells, or both cell types, which could be further skewed by distinct germline-soma ratios in testis and ovary. To distinguish these possibilities, we isolated the somatic piRNAs in the gonad, by immunoprecipitating Piwi upon germline-specific *piwi* knock-down (see methods) and then sequencing the small RNAs associated with Piwi in gonadal somatic cells (Fig. 1d). This allowed us to profile the gonadal somatic piRNA program in each of the two sexes.

Experimentally isolated gonadal somatic piRNAs from testes and ovaries (Fig. 1d) allowed us to directly compare the piRNA program in gonadal soma between sexes. We found that the *flamenco* piRNA locus shows a similar piRNA coverage profile and produces piRNAs that take up comparable fractions of total piRNAs in testicular and ovarian soma (Fig. 1e). Most of the highly expressed piRNAs in gonadal soma are antisense to transposons in both males and females, and piRNAs targeting different transposon families display a strong positive correlation between the two sexes (Pearson’s r = 0.91, *p* < 0.0001; Spearman’s r = 0.89, *p* < 0.0001; Fig. 1f). When normalized to *flamenco*, a few transposons are targeted more in either males (e.g., *idefix*) or females (e.g., *gtwin* and *mdg1*), but these biases are relatively mild (Fig. 1f). To examine piRNA production across the genome, we defined piRNA-producing loci in gonadal soma and measured their expression levels (see methods). Akin to piRNA quantification based on their transposon targets, quantifying piRNAs based on their genomic origins also revealed a strong positive correlation between the two sexes (Pearson’s r = 0.89, *p* < 0.0001; Spearman’s r = 0.82, *p* < 0.0001; Fig. 1g). We did, however, note an exception: a novel piRNA locus we identified in the gonadal soma, *77B* (Fig. 1h), produces more piRNAs in males than females (Fig. 1g). This locus resembles *flamenco*, as it makes piRNAs from one genomic strand downstream of a prominent RNA pol II peak that is indicative of a promoter, producing antisense piRNAs against transposons active in the gonadal soma (e.g., *idefix*; Fig. 1h). Nevertheless, the genome-wide view of the piRNA production in gonadal soma highly correlates between sexes. The 3’ UTR of some genes (e.g., *tj*) is known to produce piRNAs in ovarian soma^18^, and the same holds in the male counterpart (Fig. 1g). Overall, the piRNA program operating in gonadal soma shows very few sex differences.

Taking advantage of the fact that the *flamenco* piRNA locus is active exclusively in the gonadal soma but not in the germline of both testis (Extended Data Fig. 1) and ovary^6,19^, we inferred that germline piRNAs make up about 97% and 79% of the gonadal piRNAs in testis and ovary, respectively (see methods). Because germline piRNAs dominate the whole gonad piRNA pool to comparable extents in both sexes (97% / 79% = 1.2-fold difference), gonad-wide piRNA quantification is a close approximation of the germline piRNA program when studying male-female differences. These results suggest that the piRNA sexual dimorphism we observed in the whole gonads originates from the germline rather than gonadal soma. For the rest of this work, we used the gonad-wide piRNA sexual dimorphism to approximate germline piRNA sexual dimorphism, as germline piRNAs account for comparable fractions of total piRNAs in whole gonads of males and females.

### Y chromosome is necessary and sufficient to recapitulate aspects of male piRNA program

Having found that germline is the major cell type origin of piRNA sexual dimorphism, we aimed to dissect its underlying genetic control mechanisms (Fig. 2a). Distinct sex chromosome contents between sexes, specifically, the presence of Y chromosome in males, could in theory explain some sex differences in piRNAs. On the other hand, distinct sexual identities could lead to differential piRNA production even from identical piRNA loci located outside the Y. Importantly, the sex determination in *D. melanogaster* does not involve the Y^14^, which provides us with a unique opportunity to manipulate the Y chromosome without perturbing sexual identities.

**Fig. 2.**
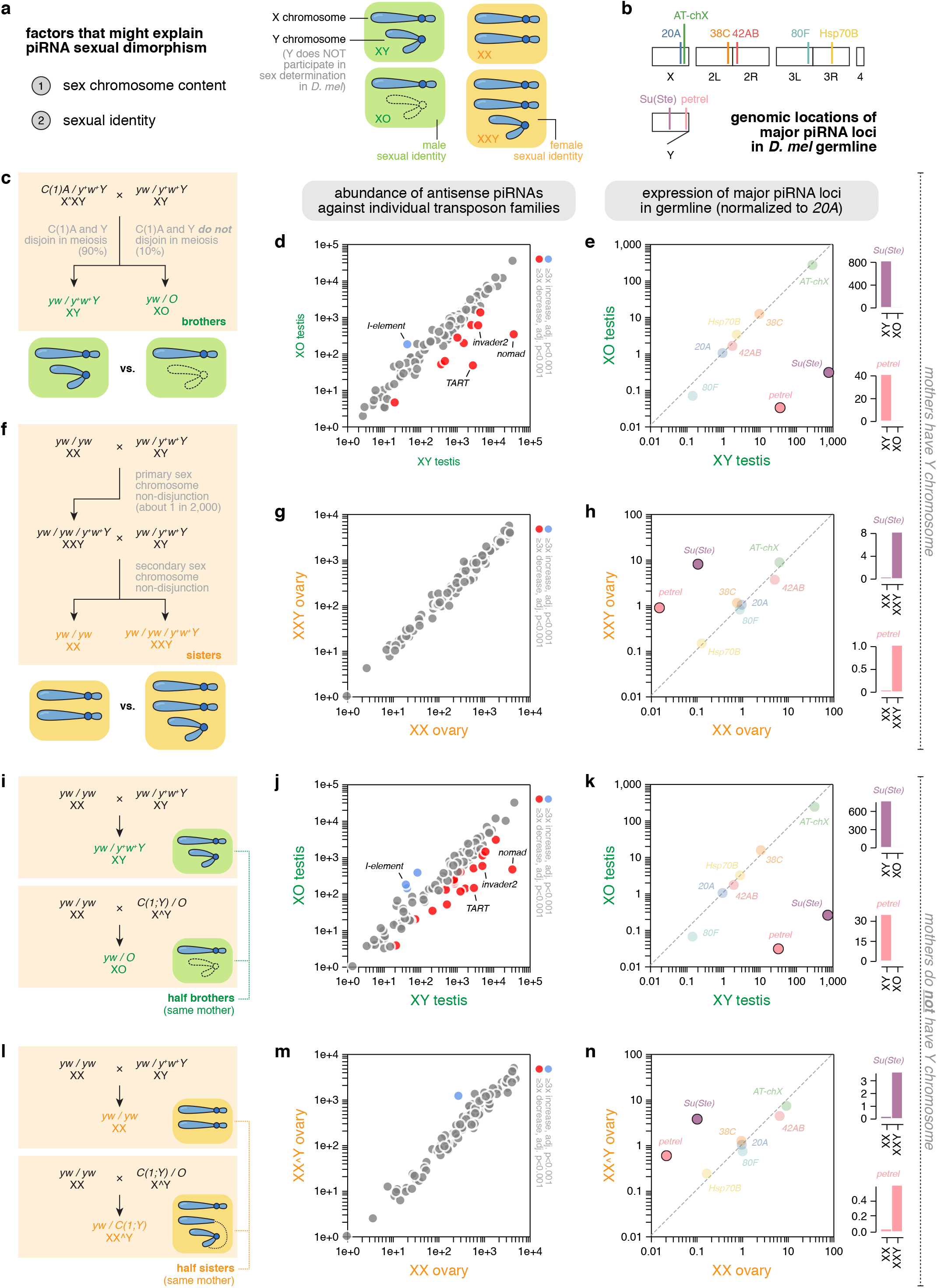
Y chromosome produces piRNAs in both males and females. **a**, Left: listed are factors that might explain piRNA sexual dimorphism. Right: cartoon showing different sex chromosome contents and respective sexual identities. X and Y chromosomes are depicted in different ways, and sexual identities are color coded with the male identity being green and the female one being orange. Note that the Y chromosome of *D. melanogaster* does not participate in sex determination, and the sex instead depends on the number of X. Hence, XY and XO are both males, and XX and XXY are both females. **b**, Illustration showing the karyotype of *D. melanogaster* with five chromosomes – X, Y, 2, 3, and 4, as well as the rough genomic locations of major piRNA loci in the germline. **c**, Cross scheme of the generation of XY and XO brothers. **d**, The abundance of transposon-targeting piRNAs in XO males compared to their XY brothers, showing the loss of piRNAs targeting several transposon families. **e**, The expression of major germline piRNA loci in XO males compared to their XY brothers, showing a specific loss of piRNAs from two Y-linked loci, *Su(Ste)* and *petrel*. **f**, Cross scheme of the generation of XX and XXY sisters. **g**, The abundance of transposon-targeting piRNAs in XXY females compared to their XX sisters, showing very limited differences. **h**, The expression of major germline piRNA loci in XXY females compared to their XX sisters, showing piRNA production from two Y-linked loci, *Su(Ste)* and *petrel*. **i**, Cross scheme of the generation of XY and XO half-brothers, with the same XX mothers. **j**, The abundance of transposon-targeting piRNAs in XO males compared to their XY half-brothers, both of which are sired by XX mothers, showing similar loss of piRNAs targeting several transposon families as seen in **d**. **k**, The expression of major germline piRNA loci in XO males compared to their XY half-brothers, both of which are sired by XX mothers, showing a similar loss of piRNAs from two Y-linked loci, *Su(Ste)* and *petrel*, as seen in **e**. **l**, Cross scheme of the generation of XX and XXY half-sisters, with the same XX mothers. **m**, The abundance of transposon-targeting piRNAs in XXY females compared to their XX half-sisters, both of which are sired by XX mothers, showing very few differences similar to that in **g**. **n**, The expression of major germline piRNA loci in XXY females compared to their XX half-sisters, both of which are sired by XX mothers, showing piRNA production from two Y-linked loci, *Su(Ste)* and *petrel*, as seen in **h**.

To pinpoint the contribution of the Y chromosome to sex differences in the piRNA program, we first generated XY and XO male sibling flies that only differ in the Y chromosome content but are otherwise genetically identical. This is done by using spontaneous sex chromosome nondisjunction that occurs at about 10% frequency in *X^XY* females carrying the compound X chromosome, *C(1)A* (Fig. 2c). When compared to their XY brothers, XO males lose piRNAs targeting several transposon families (Fig. 2d), indicating that the Y chromosome is a source of transposon-targeting piRNAs in the male. For example, the absence of the Y chromosome causes decreases in piRNAs against *nomad* and *invader2* (Fig. 2d) – two transposons that are normally targeted by more piRNAs in males than females (Fig. 1b), suggesting that these sex differences could be largely explained by the Y chromosome alone. We also note that, piRNAs targeting *I-element* appear upregulated in males lacking the Y chromosome (Fig. 2d), which warrants future investigation. Removing the Y also led to a specific loss of piRNAs from two Y-linked loci – *Su(Ste)* and *petrel* (Fig. 2b) – while leaving the piRNA production from other loci on X and autosomes unperturbed (Fig. 2b,e). Therefore, Y chromosome contributes to the male piRNA program via production of piRNAs from two loci on the Y, *Su(Ste)* and *petrel*, as well as piRNAs targeting a select group of transposons.

Complementing the male experiment, we generated XX and XXY female sibling flies that only differ in their Y chromosome contents but are otherwise genetically identical. This is achieved by first obtaining an exceptional XXY female from primary sex chromosome nondisjunction that occurs naturally about 1 in 2,000 wildtype flies^14^, and then crossing this XXY female with XY males to sire XX and XXY females through secondary sex chromosome nondisjunction (Fig. 2f). The extra Y chromosome barely altered the overall transposon-targeting piRNA program in females (Fig. 2g). Nonetheless, the presence of the Y chromosome in females triggers piRNA biogenesis from *Su(Ste)* and *petrel* (Fig. 2h), two loci that reside on the Y chromosome. Furthermore, these two Y-linked piRNA loci exhibit comparable activities to other top piRNA loci in females, including *42AB*, *38C*, and *80F* (Fig. 2h), suggesting that the Y is an active and productive piRNA source in a female cellular environment.

We generated genetically identical male and female siblings that only differ in their Y chromosome contents, however, these crosses necessitated the employment of mothers carrying a Y chromosome (Fig. 2c,f). Given that maternally deposited piRNAs instruct piRNA biogenesis in the progeny^8,10,11^, Y-bearing mothers might create a permissive environment to produce Y piRNAs in the offspring by depositing Y-derived piRNAs to the embryo. Consequently, it is unclear whether the effects of Y chromosome on male and female piRNA production we observed (Fig. 2d,e,g,h) depends on mothers carrying a Y chromosome. To empirically test the role of Y chromosome in piRNA production without mothers bearing a Y, we devised a strategy to generate half siblings of both sexes that share similar, albeit not identical, genetic backgrounds with and without Y chromosome from XX mothers (Fig. 2i,l). We observed similar effects of the Y chromosome on piRNA production when mothers do not have a Y: Y chromosome is an important source of transposon-targeting piRNAs in males but not in females (Fig. 2j,m), and the two Y-linked piRNA loci, *Su(Ste)* and *petrel,* produce piRNAs in both sexes, irrespective of the cellular sexual identity (Fig. 2k,n). We noticed that the Y exerts a slightly greater effect on the transposon-targeting piRNA program in this latter cross scheme (Fig. 2i,l) compared to the former one involving Y-bearing mothers (Fig. 2c,f), which likely results from having different fathers and thus distinct paternally inherited haploid genome. Nevertheless, the results between mothers with and without a Y chromosome are qualitatively very similar. We conclude that the presence of the Y chromosome alters the piRNA profiles independently of its presence in the mothers.

Y chromosome in *D. melanogaster* is known to exhibit imprinting effects^20–22^, that is, Y can behave differently when inherited from the mother or the father. To test if Y-linked piRNA loci show parent-of-origin effects, we designed crosses that allow females to inherit a Y chromosome from either parent. In the case of paternally inheriting the Y, we also designed crosses either with or without mothers bearing a Y (Fig. 3a top). In all cases, we detected nascent transcripts from *Su(Ste)* and *petrel* piRNA loci located on the Y chromosome (Fig. 3a middle), indicating that piRNA loci on the Y are transcriptionally active in the female germline when inherited from either parent. We also observed similar behaviors of the Y-linked piRNA loci in the male counterpart (Fig. 3b) – when inherited from either parent, with or without mothers carrying a Y, Y chromosome activates both *Su(Ste)* and *petrel* loci in the male germline. Thus, there are no obvious imprinting effects of the two Y-linked piRNA loci, and the mere presence of the Y can translate to an effect on the germline piRNA program in both sexes. Interestingly, in our cross scheme of passing the Y from mothers to daughters, *Su(Ste)* piRNA precursor transcription is also activated in the follicle cells (a gonadal somatic cell type), an unexpected finding that calls for future studies.

**Fig. 3.**
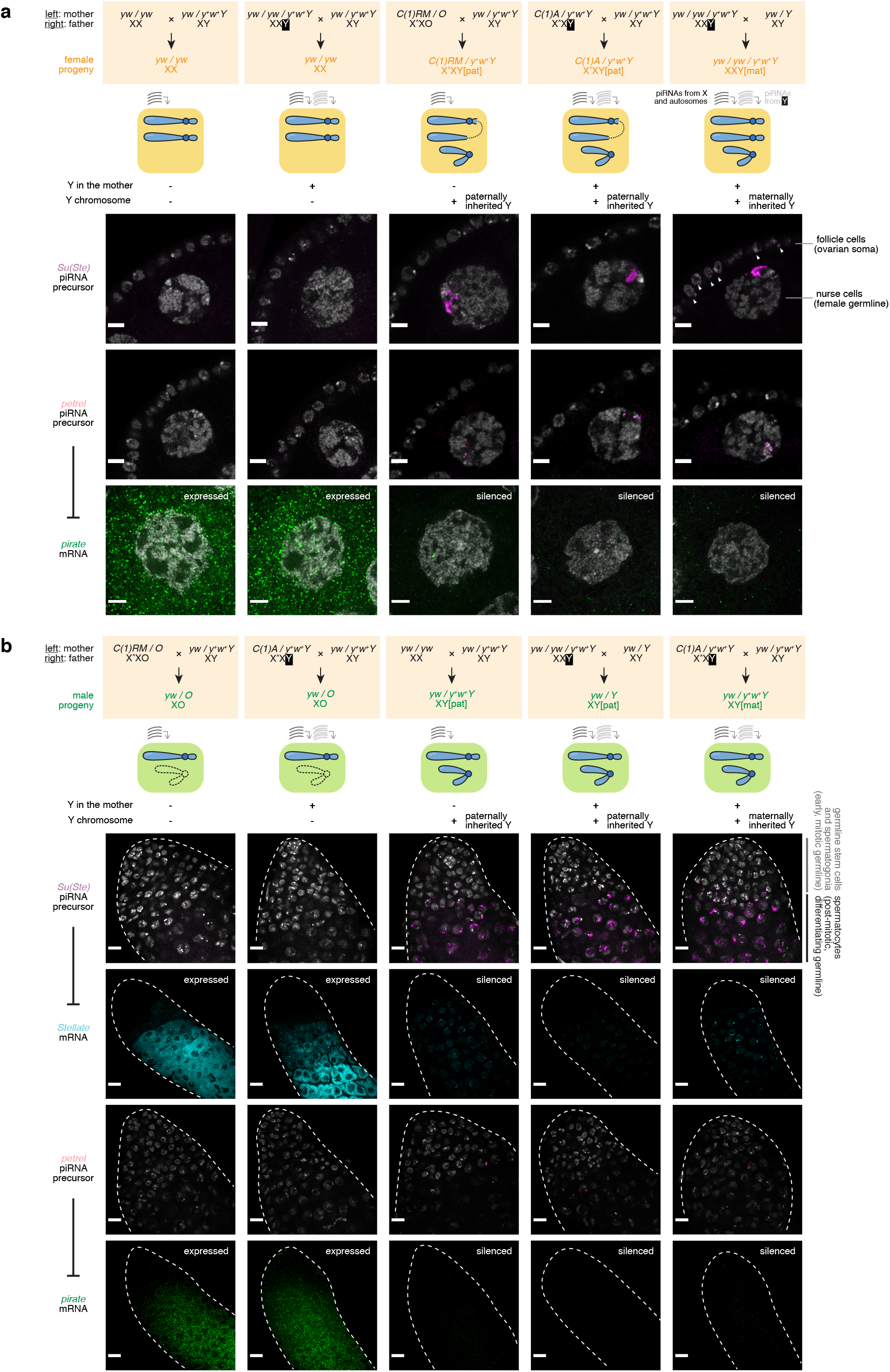
Y-linked piRNA loci are active and functional in both sexes, when inherited from either parent, regardless of whether mothers carry a Y chromosome. **a**, Top: Cross schemes that generate: XX females without mothers bearing a Y (column 1), XX females with Y-bearing mothers (column 2), XXY females without mothers bearing a Y (column 3), XXY females with Y-bearing mothers but inhering the Y from the father (column 4), and XXY females with Y-bearing mothers and inheriting the Y from the mother (column 5). Middle: cartoon showing the genotype of each kind of females generated and whether their mothers carry a Y is reflected by whether they receive maternally deposited Y-derived piRNAs. Bottom: RNA *in situ* HCR detecting *Su(Ste)* piRNA precursors (row 1), *petrel* piRNA precursors (row 2), and *pirate* mRNAs (row 3) in stage 6-7 egg chambers. Scale bar: 5μm. **b**, Top: Cross schemes that generate: XO males without mothers bearing a Y (column 1), XO males with Y-bearing mothers (column 2), XY males without mothers bearing a Y (column 3), XY males with Y-bearing mothers but inhering the Y from the father (column 4), and XY males with Y-bearing mothers and inheriting the Y from the mother (column 5). Middle: cartoon showing the genotype of each kind of males generated and whether their mothers carry a Y is reflected by whether they receive maternally deposited Y-derived piRNAs. Bottom: RNA *in situ* HCR detecting *Su(Ste)* piRNA precursors (row 1), *Stellate* mRNAs (row 2), *petrel* piRNA precursors (row 3), and *pirate* mRNAs (row 4) and at the apical tips of the testes. Scale bar: 10μm.

Whereas *Su(Ste)* piRNAs silence the *Stellate* genes that are only active in the male germline, *petrel* piRNAs silence the *pirate* gene that is ubiquitously expressed in all tissues including female germline, which allows us to explore if activating *petrel* piRNA biogenesis in the female germline leads to *pirate* silencing. In wildtype XX female germline, the *pirate* gene is active, and its transcripts can be readily detected by RNA *in situ* HCR (Fig. 3a bottom). However, introducing a Y into the female germline from either parent led to a marked silencing of the *pirate* gene (Fig. 3a bottom), suggesting that making *petrel* piRNAs in female germline has a direct functional outcome. Meanwhile, the presence of Y chromosome in mothers was neither necessary nor sufficient for the silencing of *pirate* in the female progeny. Thus, *pirate* silencing in the female germline requires the presence of the Y chromosome, regardless of the parental origin of the Y. Similarly, in the male germline, having a Y-bearing mother is neither sufficient nor required for the male germline to tame *Stellate* and *pirate*, and the presence of Y chromosome triggers silencing of both *Stellate* and *pirate* genes regardless of the Y’s inheritance path (Fig. 3b). Taken together, the differences in piRNA profiles caused by the presence of Y chromosome directly translates to differential silencing of several targets, without obvious parent-of-origin effects.

### Cellular sexual identity provides a key input into piRNA biogenesis

Though necessary, the presence of the Y chromosome in males is not sufficient to explain sex differences in the piRNA program, as piRNA loci outside the Y are also differentially expressed in two sexes (Fig. 1c). What underlies piRNA sexual dimorphism outside the Y? In *D. melanogaster*, a cascade of molecular switches takes place after counting the number of X, culminating in either male or female cellular sexual identity^14–16,23–28^ (Fig. 4a). To examine the contribution of sexual identities to germline piRNA sexual dimorphism without confounding impacts of the Y chromosome, we sought to masculinize XX female germline and compare it to XO male germline that lacks a Y (Fig. 4b). Unlike the *Drosophila* soma, where sex determination occurs cell-autonomously, *Drosophila* germline receives an additional input from the soma on top of its own chromosomal content to determine the germline sex^24,25^. When the germline sex does not match the somatic sex, germline either dies or becomes tumorous^29,30,23–25,31,32^, so a productive germline sex reversal requires perturbing both the germline and somatic sex. Given that there is very little sexual dimorphism in the gonadal somatic piRNAs (Fig. 1e,f,g,h), reversing the somatic sex should not confound our study of germline piRNA sexual dimorphism. Hence, our germline sex reversal was done in sex-reversed soma, which allowed us to interrogate the effect of sexual identities on germline piRNA sexual dimorphism.

**Fig. 4.**
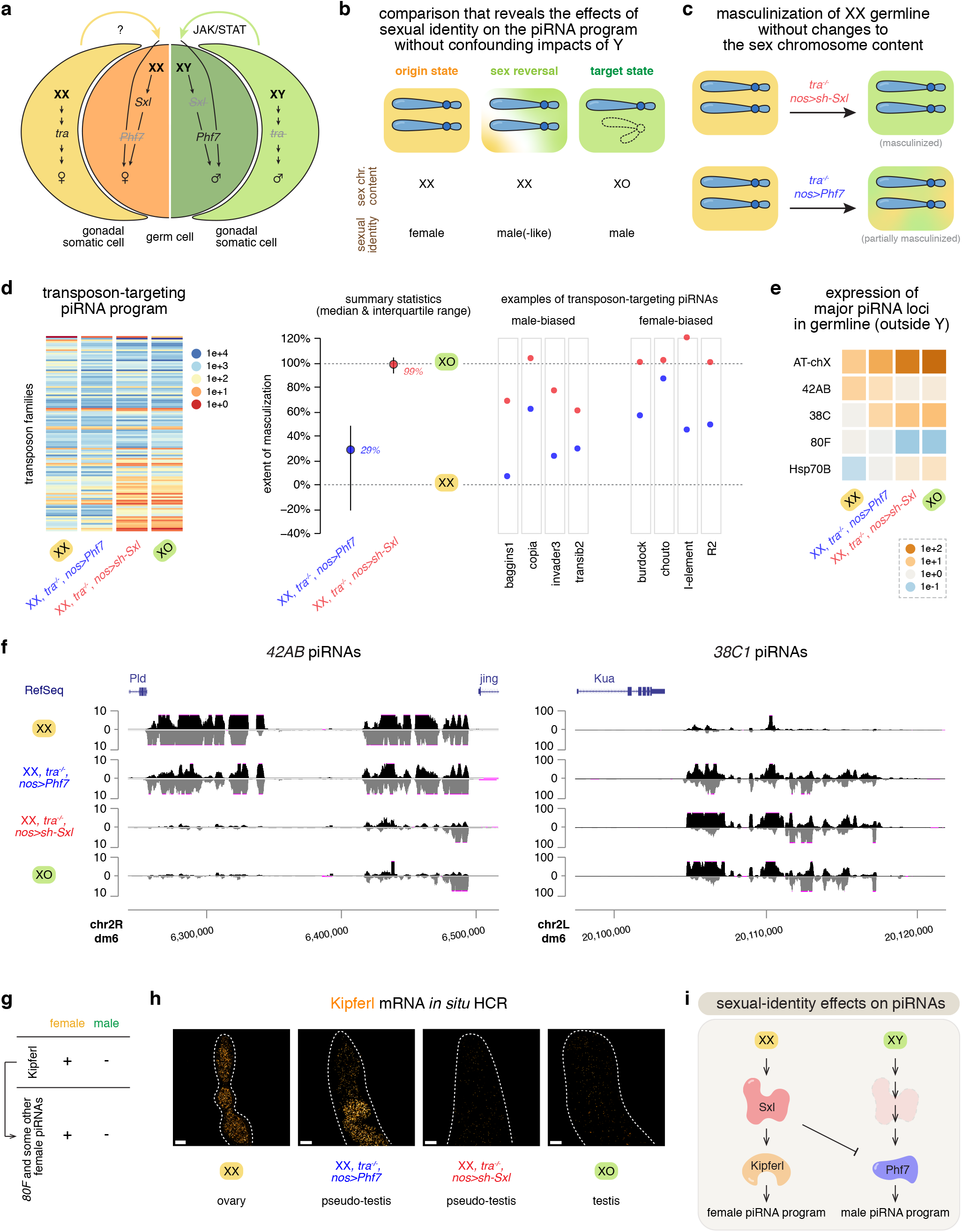
Sexual identity provides a key input into piRNA biogenesis and is a major determinant of the piRNA program. **a**, A simplified model of the sex determination pathway in germline and soma. On top of its own chromosome content, germline receives an additional input from the soma to determine its sex. **b**, A comparison scheme that uses sex reversal to examine the effects of sexual identities on the piRNA program. XX female germline is masculinized and compared to both wildtype XX females (the origin state) and XO males (the target state). Thus, any differences observed would reveal the effects of sexual identities on piRNAs, without confounding impacts of the Y chromosome. **c**, Cartoon showing the masculinization of the XX female germline by genetic perturbations, without any changes to the sex chromosome content. To facilitate germline masculinization, the soma is masculinized by mutating *tra*. In addition, germline-specific knock-down of *Sxl* near completely masculinizes the female germline, while ectopic expression of *Phf7* in the germline led to partial masculinization. **d**, Left: a heatmap showing the abundance of piRNAs targeting different transposons in XX female, XO male, or XX masculinized by perturbing either *Phf7* or *Sxl* expression (in the *tra* mutant background). Each row represents piRNAs that target a different transposon, and their expression levels are color coded. Middle: Quantification of the extent of masculinization for piRNAs targeting individual transposon families. For each transposon family, the abundance of corresponding antisense piRNAs in XX female is scaled to 0% and that in XO male is scaled to 100%, so the levels of transposon-targeting piRNAs in masculinized XX can be normalized to reflect the extent of masculinization. Shown are the summary statistics (median and interquartile range) of the antisense piRNAs targeting different transposon families. Right: piRNA abundance upon XX masculinization for four examples of male-biased and female-biased transposon-targeting piRNAs, respectively. In these examples as well as the overall summary statistics, perturbing *Sxl* led to a stronger masculinization of the piRNA program than perturbing *Phf7*. **e**, A heatmap showing the expression of major germline piRNA loci (located outside Y) in XX female, XO male, or XX masculinized by perturbing either *Phf7* or *Sxl* expression (in the *tra* mutant background). **f**, UCSC genome browser view of the piRNA coverage profiles over the locus *42AB* (left) and the locus *38C1* (right) in XX female, XO male, or XX masculinized by perturbing either *Phf7* or *Sxl* expression (in the *tra* mutant background). **g**, Female-specific expression of Kipferl and Kipferl-dependent piRNAs. **h**, RNA *in situ* HCR detecting *Kipferl* mRNA in XX female, XO male, or XX masculinized by perturbing either *Phf7* or *Sxl* expression (in the *tra* mutant background). **i**. A genetic circuit that connects the sex determination pathway and piRNA biogenesis.

To explore if and how sexual identities impact germline piRNA profiles, we masculinized XX female germline by germline-specific knock-down of *Sex lethal* (*Sxl*), the major factor that governs the female identity in germline^23–26^, in the *transformer* (*tra*) mutant background that has a masculinized soma^29^ (Fig. 4c). Strikingly, masculinizing the XX female germline converted its transposon-targeting piRNA program to a state that closely resembles the XO male germline (Fig. 4d left). When quantified, the median extent of masculinization for piRNAs targeting different transposon families is 99% (see methods; Fig. 4d right). Similarly, for major piRNA loci outside the Y, their expression levels and piRNA coverage profiles also switched from an XX female state to an XO male state upon masculinization of the XX germline (Fig. 4e,f). For example, abundant piRNAs are made from both proximal and distal ends of the *42AB* piRNA locus in XX females, but masculinized XX germline only make some piRNAs from the distal end of *42AB* and barely any from the proximal side, reminiscent of the XO male germline (Fig. 4f left). In sum, reversing the germline sexual identity is sufficient to switch the germline piRNA program from one sex to the other, suggesting that the cellular sexual identity provides a key input into piRNA biogenesis.

How does the germline interpret its sexual identity to elicit a sex-specific piRNA program? We showed that this sexual-identity effect on piRNA biogenesis is governed by the major switch protein Sxl (Fig. 4d,e,f), which is active in the female, but not male, germline. Next, we looked into how Sxl orchestrates a female-specific piRNA program in the germline. Sxl is known to regulate two target genes that exhibit sex-specific expression patterns in the germline^33^: Tdrd5l^28^, a cytoplasmic protein that forms granules distinct from the piRNA processing sites, and Phf7^34^, a chromatin reader protein that binds H3K4me2/3. Both Tdrd5l and Phf7 promote a male identity, and Sxl represses these two factors to bolster a female identity. Since Tdrd5l and Phf7 act genetically redundantly to support a male identity^28^, we focused on Phf7 for this study and asked whether and, if so, to what extent Phf7 mediates the sexual-identity effect on piRNA biogenesis. Expressing Phf7 in the female germline accompanied by somatic masculinization partially masculinized the XX female germline, leading to a 29% median extent of masculinization of transposon-targeting piRNAs (Fig. 4d). For many transposon-targeting piRNAs (e.g., those targeting *copia* and *burdock*), the ectopic expression of Phf7 in XX germline shifted the piRNA profile from a female state towards a male state, but not as completely as losing Sxl did (Fig. 4d right). This partial reversal of the piRNA program from one sex to the other by Phf7 activation is also obvious when examining the expression of major piRNA loci in the genome. Each of the major piRNA loci in Phf7-expressing XX female germline resumed an activity somewhere in between the wildtype XX female and XO male, for both male- and female-biased loci (Fig. 4e,f). For instance, Phf7 dampened the activity of *42AB*, a female-biased piRNA locus, and enhanced the activity of *38C*, a male-biased piRNA locus (Fig. 4e,f). These observations indicate that Phf7 promotes a male piRNA program, and Sxl supports a female piRNA program in part through repressing Phf7. Thus, Phf7 mediates part of the sexual-identity effect on piRNA biogenesis, acting downstream of Sxl.

Recently, a female-specific piRNA biogenesis factor, the zinc-finger protein Kipferl, was described to drive a subset of piRNA production in the female germline^35^. In particular, piRNA production from *80F*, a sex-specific piRNA locus only active in the female germline (Fig. 4e), depends on Kipferl^35^ (Fig. 4g), indicating that Kipferl is directly responsible for some of the germline piRNA sexual dimorphism. As Kipferl appears dedicated to the piRNA pathway and is absent in the male germline, we hypothesized that Kipferl is an effector protein that acts downstream of the sex determination pathway to elaborate a female piRNA program. Indeed, knocking-down *Sxl* in the female germline is sufficient to abrogate the expression of *Kipferl* in XX germline, suggesting that Kipferl expression depends on Sxl (Fig. 4h). On the other hand, expressing Phf7 in the XX female germline did not perturb *Kipferl* expression (Fig. 4h), suggesting that Phf7 and Kipferl act in parallel, both downstream of Sxl, to promote male and female piRNA programs, respectively (Fig. 4i).

Taken together, we elucidated a genetic circuit that connects the sex determination pathway to germline piRNA sexual dimorphism (Fig. 4i). XX germline activates Sxl, which positively regulates Kipferl to produce female-specific piRNAs and negatively regulates Phf7 to suppress a male piRNA program. On the contrary, XY germline lacks functional Sxl to activate Kipferl and instead expresses Phf7 to elaborate a male piRNA program. We conclude that male and female sexual identities enable divergent piRNA production programs, sculpting a sexually dimorphic molecular trait alongside the male-specific Y chromosome.

## Discussion

In this work, we identified the germline as the source of piRNA sexual dimorphism in fly gonads. Building on this, we deciphered the genetic control underlying the sex-specific piRNA program. We characterized the contribution of the Y chromosome to the male piRNA program, and we showed that the presence of the Y is sufficient to recapitulate some aspects of the male piRNA program in a female cellular environment. In fact, the effect of the Y is independent of its parental origin and mothers’ sex chromosome contents. The ability of Y-linked piRNA loci to act in both male and female cellular environments independently of its inheritance path implies unique regulatory mechanisms^36^ employed by the Y and distinctive evolutionary forces acting on the Y. Meanwhile, we showed that sexual identity is another major determinant of the piRNA program that regulates piRNA biogenesis outside the Y. Specifically, sexual identity shapes piRNA sexual dimorphism under the control of Sxl, which relays the sexual identity of a cell to piRNA biogenesis through the histone reader protein Phf7 and the zinc-finger protein Kipferl. We speculate that the sex determination pathway has been hijacked by transposons to facilitate their sex-biased germline invasion^12^, so integrating the information of germline sexual identities into piRNA biogenesis provides a means to directly couple the sex-specific piRNA defense program with sex-specific transposon threats. Together, our work revealed that sex chromosome and sexual identity control different facets of piRNA sexual dimorphism, and it is their collective action that sculpts the sex-specific piRNA program in fly germline. It is very likely that other sexually dimorphic traits are under the control of both sex chromosome and sexual identity, and disentangling the effects of the two promises to offer new insights into how a molecular pathway can be modified by each of the two sexes to execute essential functions.

## Acknowledgement

We thank Grace YC Lee, Felipe Karam Teixeira, Justin Blumenstiel, Katalin Fejes Toth, and members of the Aravin Laboratory for discussion, Jim Kennison for advice on sex chromosome nondisjunction, Mark Van Doren and Helen Salz for advice on sex determination, and Yukiko Yamashita for sharing unpublished results. We thank Angela Stathopoulos, Ellen Rothenberg, and Henry Chung for comments on the manuscript. We are grateful to Liz Gavis, Bloomington Drosophila Stock Center and Vienna Drosophila Resource Center for fly lines. We appreciate the help of Igor Antoshechkin (Millard and Muriel Jacobs Genetics and Genomics Laboratory, Caltech) with sequencing, the help of Fan Gao (Bioinformatics Resource Center, Caltech) with bioinformatic analysis, the help of Grace Shin (Molecular Technologies, Caltech) with *in situ* HCR, and the help of Giada Spigolon and Andres Collazo (Biological Imaging Facility, Caltech) with microscopy. This work was supported by grants from the National Institutes of Health (R01 GM097363) and by the HHMI Faculty Scholar Award to A.A.A.

---

**Extended Data Fig. 1.**
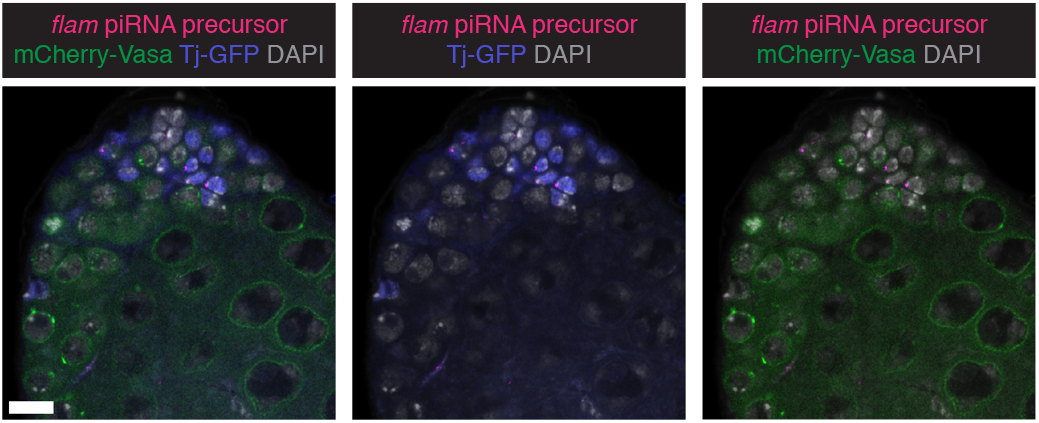
The *flamenco* piRNA locus is only active in the gonadal soma in testis. RNA *in situ* HCR detecting *flamenco* piRNA precursors in testes expressing mCherry-Vasa and Tj-GFP under endogenous regulatory elements, stained for DAPI. *flamenco* transcripts are only detected in gonadal somatic cells (including both early cyst cells marked by Tj expression and hub cells marked by the lack of Tj and Vasa expression), but not in germline cells (marked by Vasa expression). Scale bar: 10μm.

## Methods

### Fly stocks and crosses

Stocks and crosses were raised at 25 °C. The following stocks were used: *yw* (BDSC 6599), *y^+^w^+^*Y (BDSC 7060), *C(1)A* (BDSC 2950), *C(1)RM* (BDSC 9460), *C(1;Y)* (BDSC 9460)*, nos-Gal4* (BDSC 4937), *bam-Gal4* (BDSC 80579), *sh-piwi* (BSDC 33724), *sh-Sxl* (BDSC 38195), *UAS-Phf7* (BDSC 15894), *tra*^*1*^ (BDSC 675) and *Df(tra)* (BDSC 5416) were obtained from Bloomington Drosophila Stock Center, and *Tj-GFP* (VDRC 318066) was obtained from Vienna Drosophila Resource Center. *GFP-Piwi* (BAC) and *mCherry-Vasa* were previously described^37,38^. To minimize genetic background differences, *yw / y^+^w^+^*Y was backcrossed to the inbred *yw* line for six consecutive generations, via a single male at every generation. Similarly, after generating *C(1)A / y^+^w^+^*Y females, we backcrossed them to *yw / y^+^w^+^*Y males for six consecutive generations, via 2-3 females at every generation. To obtain an XXY exceptional female, we looked for a female carrying the marked Y chromosome (*y^+^w^+^*Y) in the *yw / y^+^w^+^*Y stock, which typically took no more than two months. To deplete germline Piwi, we expressed *sh-piwi* using both *nos-Gal4* and *bam-Gal4,* which led to efficient knock-down of Piwi in the germline as evidenced by the loss of germline GFP-Piwi expression in both testis and ovary. For sex reversal experiments, a Y chromosome marked by *B*^*S*^ (present in the *Df(tra)* stock) that alters the eye shape was employed, such that the sex chromosome content could be inferred independently of the morphological sex.

### RNA *in situ* hybridization and RNA *in situ* hybridization chain reaction (HCR)

For RNA *in situ* HCR, probes, amplifiers and buffers were purchased from Molecular Instruments (molecularinstruments.org) for *flam* (3893/E046), *petrel* (3872/E024), *pirate* (3916/E064), *Stellate* (4537/E832) and *Kipferl* (4708/E1062) transcripts. RNA *in situ* HCR v3.0^39^ was done according to manufacturer’s recommendations for generic samples in solution. To detect *Su(Ste)* transcripts, we did conventional RNA *in situ* hybridization using DNA probes (75bp, position 994-1068 of *Su(Ste): CR42424*, sense direction) directly conjugated with fluorophore purchased from IDT.

### Image acquisition and processing

Confocal images were acquired with Zeiss LSM 800 or LSM 980 using a 63x oil immersion objective (NA=1.4) and processed using Fiji^40^. Single focal planes were shown in all images, where dotted outlines were manually drawn for illustration purposes.

### Small RNA sequencing

Argonaute-associated small RNAs were isolated from ovaries (20 pairs per sample) or testes (30 pairs per sample) using TraPR columns^41^. Purified small RNA was subject to library prep using NEBNext Multiplex Small RNA Sample Prep Set for Illumina (NEB E7330). Adaptor-ligated, reverse-transcribed, PCR-amplified samples were purified again by PAGE (6% polyacrylamide gel), from which we cut out the band within the desired size range. This additional size selection by PAGE eliminated other, longer RNAs (>30 nt) captured by TraPR columns. To isolate Piwi-associated small RNAs in gonadal soma, we first immunoprecipitated GFP-Piwi from gonads lacking germline Piwi (see above for fly crosses) using GFP-Trap (ChromoTek) magnetic agarose beads, as described before^42^. Small RNAs associated with gonadal somatic Piwi are then purified by TraPR columns and library-prepared, as described above for all Argonaute-associated small RNAs. Two biological replicates per genotype were sequenced on Illumina HiSeq 2500.

### Analysis of transposon-targeting piRNAs

To computationally extract piRNAs from all Argonaute-associated small RNAs, adaptor-trimmed small RNAs were size-selected for 23-29nt (cutadapt 2.5) and those mapped to rRNA, tRNA, snRNA, snoRNA, miRNA, hpRNA and 7SL RNA were discarded (bowtie 1.2.2 with -v 3). piRNAs were then mapped to RepBase25.08 (manually curated) and those antisense to transposon consensus sequences with ≤3 mismatches are designated as transposon-targeting piRNAs. For LTR transposons, reads mapping to the LTR and internal sequences of a given transposon family were merged for quantification, given their well-correlated behaviors. All quantification was done using the mean of two biological replicates.

### Analysis of the expression of major piRNA loci

piRNAs were computationally extracted as described above. piRNAs were mapped to the dm6 genome using a previously described algorithm^12^ that considers both piRNAs that map to single unique positions in the genome as well as those that map to “local repeats” (defined as repeats that are contained within a genomic window <2Mb in length in dm6 reference genome). Major piRNA loci, their coordinates and quantification method were described before^12^. The average of two biological replicates was shown in all figures.

### Definition of piRNA-producing loci in gonadal soma

piRNA-producing loci in gonadal soma were defined as previously described for piRNA-producing loci (also known as “piRNA clusters”) in whole gonads^12^. Briefly, piRNAs isolated from gonadal soma were mapped to the genome and those that map uniquely or to local repeats were kept and quantified over 1Kb windows that tile the entire genome. Neighboring 1Kb widows within 3Kb were merged. If merged windows were ≥5Kb, they were merged again within 15Kb, and this process was repeated twice. This *de novo* method recapitulated the *flamenco* locus and the 3’UTR of the *tj* gene – two loci that are known to make abundant piRNAs in ovarian soma – confirming its utility.

### Inference of germline contribution to whole gonad piRNAs

Given that *flamenco* is only active in the gonadal soma but not in the germline, *flamenco* piRNAs found in whole gonad piRNAs must come from somatic cells in the gonad. Experimentally isolated gonadal somatic piRNAs revealed the contribution of *flamenco* piRNAs to total piRNAs in the gonadal soma (e.g., 25%), so if *flamenco* takes up 5% of whole gonad piRNAs, gonadal soma will contribute to 20% (5% / 25% = 20%) of whole gonad piRNAs. Then, the germline contribution to whole gonad piRNAs is 100% - 20% = 80%. When calculating actual contributions of *flamenco* piRNAs to gonadal soma and whole gonads of both sexes, the mean of two replicates was used.

### Data visualization and statistical analysis

All data visualization and statistical analysis were done in Python 3 via JupyterLab with the following software packages: numpy^43^, pandas^44^ and altair^45^. The UCSC Genome Brower^46^ and IGV^47,48^ were used to explore sequencing data and to prepare genome browser track panels shown. Two biological replicates showed similar coverage profiles on the genome browser, so one of the two replicates was randomly selected to be shown in the figures.

### Data availability

Sequencing data will be uploaded to NCBI SRA.

## References

1. Kopp, A., Duncan, I., Godt, D. & Carroll, S. B. Genetic control and evolution of sexually dimorphic characters in Drosophila. Nature 408, 553–559 (2000).

2. Williams, T. M. et al. The regulation and evolution of a genetic switch controlling sexually dimorphic traits in Drosophila. Cell 134, 610–623 (2008).

3. Cox, R. M. & Calsbeek, R. Sexually antagonistic selection, sexual dimorphism, and the resolution of intralocus sexual conflict. Am Nat 173, 176–187 (2009).

4. Galouzis, C. C. & Prud’homme, B. Transvection regulates the sex-biased expression of a fly X-linked gene. Science 371, 396–400 (2021).

5. Vagin, V. V. et al. A distinct small RNA pathway silences selfish genetic elements in the germline. Science 313, 320–324 (2006).

6. Brennecke, J. et al. Discrete Small RNA-Generating Loci as Master Regulators of Transposon Activity in Drosophila. Cell 128, 1089–1103 (2007).

7. Saito, K. et al. Specific association of Piwi with rasiRNAs derived from retrotransposon and heterochromatic regions in the Drosophila genome. Genes Dev. 20, 2214–2222 (2006).

8. Brennecke, J. et al. An epigenetic role for maternally inherited piRNAs in transposon silencing. Science 322, 1387–1392 (2008).

9. Malone, C. D. et al. Specialized piRNA Pathways Act in Germline and Somatic Tissues of the Drosophila Ovary. Cell 137, 522–535 (2009).

10. de Vanssay, A. et al. Paramutation in Drosophila linked to emergence of a piRNA-producing locus. Nature 490, 112–115 (2012).

11. Le Thomas, A. et al. Transgenerationally inherited piRNAs trigger piRNA biogenesis by changing the chromatin of piRNA clusters and inducing precursor processing. Genes Dev. 28, 1667–1680 (2014).

12. Chen, P. et al. piRNA-mediated gene regulation and adaptation to sex-specific transposon expression in D. melanogaster male germline. Genes Dev 35, 914–935 (2021).

13. Chen, P., Luo, Y. & Aravin, A. A. RDC complex executes a dynamic piRNA program during Drosophila spermatogenesis to safeguard male fertility. PLoS Genet 17, e1009591 (2021).

14. Bridges, C. B. Non-Disjunction as Proof of the Chromosome Theory of Heredity. Genetics 1, 1–52 (1916).

15. Bridges, C. B. Triploid Intersexes in Drosophila melanogaster. Science 54, 252–254 (1921).

16. Erickson, J. W. & Quintero, J. J. Indirect effects of ploidy suggest X chromosome dose, not the X:A ratio, signals sex in Drosophila. PLoS Biol 5, e332 (2007).

17. Li, C. et al. Collapse of Germline piRNAs in the Absence of Argonaute3 Reveals Somatic piRNAs in Flies. Cell 137, 509–521 (2009).

18. Robine, N. et al. A broadly conserved pathway generates 3’UTR-directed primary piRNAs. Curr Biol 19, 2066–2076 (2009).

19. Pélisson, A. et al. Gypsy transposition correlates with the production of a retroviral envelope-like protein under the tissue-specific control of the Drosophila flamenco gene. EMBO J 13, 4401–4411 (1994).

20. Maggert, K. A. & Golic, K. G. The Y chromosome of Drosophila melanogaster exhibits chromosome-wide imprinting. Genetics 162, 1245–1258 (2002).

21. Menon, D. U. & Meller, V. H. Imprinting of the Y chromosome influences dosage compensation in roX1 roX2 Drosophila melanogaster. Genetics 183, 811–820 (2009).

22. Lemos, B., Branco, A. T., Jiang, P.-P., Hartl, D. L. & Meiklejohn, C. D. Genome-wide gene expression effects of sex chromosome imprinting in Drosophila. G3 (Bethesda) 4, 1–10 (2014).

23. Schüpbach, T. Normal female germ cell differentiation requires the female X chromosome to autosome ratio and expression of sex-lethal in Drosophila melanogaster. Genetics 109, 529–548 (1985).

24. Steinmann-Zwicky, M., Schmid, H. & Nöthiger, R. Cell-autonomous and inductive signals can determine the sex of the germ line of drosophila by regulating the gene Sxl. Cell 57, 157–166 (1989).

25. Nöthiger, R., Jonglez, M., Leuthold, M., Meier-Gerschwiler, P. & Weber, T. Sex determination in the germ line of Drosophila depends on genetic signals and inductive somatic factors. Development 107, 505–518 (1989).

26. Hashiyama, K., Hayashi, Y. & Kobayashi, S. Drosophila Sex lethal gene initiates female development in germline progenitors. Science 333, 885–888 (2011).

27. Shapiro-Kulnane, L., Smolko, A. E. & Salz, H. K. Maintenance of Drosophila germline stem cell sexual identity in oogenesis and tumorigenesis. Development 142, 1073–1082 (2015).

28. Primus, S., Pozmanter, C., Baxter, K. & Van Doren, M. Tudor-domain containing protein 5-like promotes male sexual identity in the Drosophila germline and is repressed in females by Sex lethal. PLoS Genet 15, e1007617 (2019).

29. Sturtevant, A. H. A Gene in Drosophila Melanogaster That Transforms Females into Males. Genetics 30, 297–299 (1945).

30. Brown, E. H. & King, R. C. Studies on the Expression of the Transformer Gene of Drosophila Melanogaster. Genetics 46, 143–156 (1961).

31. Steinmann-Zwicky, M. Sex determination in Drosophila: sis-b, a major numerator element of the X:A ratio in the soma, does not contribute to the X:A ratio in the germ line. Development 117, 763–767 (1993).

32. Casper, A. & Van Doren, M. The control of sexual identity in the Drosophila germline. Development 133, 2783–2791 (2006).

33. Grmai, L., Pozmanter, C. & Van Doren, M. The Regulation of Germline Sex Determination in Drosophila by Sex lethal. Sex Dev 1–6 (2022) doi:10.1159/000521235.

34. Yang, S. Y., Baxter, E. M. & Van Doren, M. Phf7 controls male sex determination in the Drosophila germline. Dev Cell 22, 1041–1051 (2012).

35. Baumgartner, L. et al. The Drosophila ZAD zinc finger protein Kipferl guides Rhino to piRNA clusters. Elife 11, e80067 (2022).

36. Venkei, Z. G. et al. Drosophila Males Use 5′-to-3′ Phased Biogenesis to Make Stellate-silencing piRNAs that Lack Homology to Maternally Deposited piRNA Guides. https://www.biorxiv.org/content/10.1101/2022.09.12.507655v1.full (2022) doi:10.1101/2022.09.12.507655.

37. Le Thomas, A. et al. Piwi induces piRNA-guided transcriptional silencing and establishment of a repressive chromatin state. Genes Dev. 27, 390–399 (2013).

38. Lerit, D. A. & Gavis, E. R. Transport of germ plasm on astral microtubules directs germ cell development in Drosophila. Curr Biol 21, 439–448 (2011).

39. Choi, H. M. T. et al. Third-generation in situ hybridization chain reaction: multiplexed, quantitative, sensitive, versatile, robust. Development 145, (2018).

40. Schindelin, J. et al. Fiji: an open-source platform for biological-image analysis. Nat. Methods 9, 676–682 (2012).

41. Grentzinger, T. et al. A universal method for the rapid isolation of all known classes of functional silencing small RNAs. Nucleic Acids Res 48, e79 (2020).

42. Ninova, M. et al. Su(var)2-10 and the SUMO Pathway Link piRNA-Guided Target Recognition to Chromatin Silencing. Mol Cell 77, 556–570.e6 (2020).

43. Oliphant, T. E. Guide to NumPy. (Continuum Press, 2015).

44. McKinney, W. Data Structures for Statistical Computing in Python. in 56–61 (2010). doi:10.25080/Majora-92bf1922-00a.

45. VanderPlas, J. et al. Altair: Interactive Statistical Visualizations for Python. JOSS 3, 1057 (2018).

46. Kent, W. J. et al. The human genome browser at UCSC. Genome Res. 12, 996–1006 (2002).

47. Robinson, J. T. et al. Integrative genomics viewer. Nat Biotechnol 29, 24–26 (2011).

48. Thorvaldsdóttir, H., Robinson, J. T. & Mesirov, J. P. Integrative Genomics Viewer (IGV): high-performance genomics data visualization and exploration. Brief. Bioinformatics 14, 178–192 (2013).

